# Causal assessment of gene regulatory network in single-cell transcriptomics data based on Bayesian networks

**DOI:** 10.64898/2025.12.17.695014

**Authors:** Noriaki Sato, Marco Scutari, Seiya Imoto

## Abstract

Gene regulatory network (GRN) inference is an essential tool for revealing dysregulated relationships between genes in different cell types from single-cell transcriptomic (SCT) data. GRNs based on Bayesian networks (BNs) learned from SCT data can elucidate directed regulatory relationships representing complex disease mechanisms and their interplay through graphical modeling. However, software for learning BNs from SCT data is not widely available, nor is evaluating the BNs’ structural accuracy in representing causal relationships between genes. Here, we describe the scstruc R package. This package provides a suite of BN structure learning algorithms specifically designed for handling SCT data; evaluating the resulting networks based on the causal relationships they represent regardless of the availability of established molecular interaction networks; and comparing regulatory relationships between conditions. We demonstrated that scstruc can identify biologically relevant differential regulatory relationships between groups on a per-cell basis. The package is available at https://github.com/noriakis/scstruc.

## Motivation

Gene regulatory network inference from single-cell transcriptomic data offers valuable insights into dysregulated gene interactions at cellular resolution. While existing studies on inference methods typically evaluate performance using predictive metrics such as the area under the precision-recall curve, our understanding remains limited regarding how these algorithms perform in capturing causal relationships between genes. Moreover, there is a lack of frameworks to assess such performance effectively.

## Introduction

Single-cell transcriptomics (SCT) is widely used in biology and medicine to develop fundamental insights into disease mechanisms.^1,2^ The analysis of SCT data involves many tasks, such as cell clustering and annotation,^3,4^ supporting the inference of gene regulatory networks (GRN). Among the software tools that implement them,^5^ two options are *SCENIC*, which uses transcription factor and gene relationships,^6^ and *hdWGCNA,* which uses the weighted gene correlation network analysis to evaluate GRNs learned from SCT data.^7^

Bayesian networks (BNs) are a class of graphical models defined over a set of random variables corresponding to the nodes of a directed acyclic graph (DAG). BNs have been used in many studies to infer GRNs from transcriptomic data, measuring mRNA collected using microarray or bulk RNA-seq.^8–10^ Unlike undirected network models, directed graphs assign directions to the regulatory relationships between genes. BNs allow for probabilistic reasoning based on the structure learned from interventional data, observational data, or their combination, which may be available depending on the cost of perturbation experiments.^11–15^ Under additional assumptions, such as causal sufficiency, we can use BNs as causal networks and perform causal reasoning by simulating interventions and counterfactuals.^16,17^ While there is software to estimate directed relationships from time-series expression data,^18^ there are few alternatives for learning directed GRNs from the observational, cross-sectional data that are integrated into existing SCT data analysis pipelines.

We can use several machine learning approaches to learn GRNs based on BNs from data. Han et al. successfully used penalized regressions for learning GRNs as sparse Gaussian BNs from ovarian adenocarcinoma tumor samples and the mRNA profiles in the Cancer Cell Line Encyclopedia.^19^ Xu et al. used multi-omics integration based on BNs to reveal the mechanisms of insulin resistance.^20^ Choi and Ni have used BNs building on the zero-inflated Poisson model to identify key hub genes in primary colorectal cancer tumors.^21^ As an alternative, BNs based on the hurdle model build on earlier undirected networks for zero-inflated data.^22,23^ Also, the use of variational autoencoders for jointly modeling GRN and gene expression in SCT data, generative adversarial network, and the gradient boosting-based inference for learning neighbors of each gene from the data has been proposed.^24–26^

While these studies have investigated the performance of GRN inference and the ability of GRNs based on machine learning approaches to answer specific clinical questions, few have structurally evaluated these GRNs as mechanistic systems models encoding causal relationships. Networks that faithfully reproduce actual biological causal relationships are ideal tools for hypothesis generation because the propagation of the effects of simulated perturbations through the network closely mirrors the impact of the corresponding molecular interventions.^27,28^ The causal discovery literature has proposed many structural metrics for evaluating BNs,^29–31^ but they have not found applications to GRNs because of a combination of high computational costs, inconsistencies in how they score causally-equivalent networks learned from observational data, and poor alignment with established, practical quality metrics for molecular networks.

These theoretical and methodological limitations are compounded by the limited availability of software that can learn BNs from SCT data and evaluate their structure in a probabilistically-correct, reliable, scalable, and user-friendly way. To address this need, we have developed the R package *scstruc,* which we will describe in the remainder of this paper. Its main features are:

● Adaptations of several structure learning algorithms to SCT data based on widely used suitable data structures such as *SingleCellExperiment*.^4^
● Systematic evaluation of learned networks in terms of the causal relationships they encode by comparing multiple methods and without assuming the availability of a reference ground-truth network. Standard biological interaction databases are used for further validation.
● Differential identification of regulatory pathways between different groups.

*scstruc* aims to facilitate and streamline the application of BNs to SCT data. In turn, the ability to learn GRNs will help generate novel hypotheses to drive biological and clinical experiments and provide a deeper understanding of the mechanisms underlying SCT data.

## Results

### scstruc architecture

*scstruc* is an R package that infers directed GRNs and evaluates the resulting networks based on their causal validity, both in situations where the reference networks are available or not. The workflow of GRN inference and evaluation using *scstruc* is illustrated in Figure 1. The first step is to perform structure learning based on various algorithms tailored for the nature of SCT data. Following the structure learning, the networks are compared against ground-truth directed networks. The ground-truth network can be constructed based on the user’s biological pathway of interest using internal functions of the software. Furthermore, the package implements the Intersection-Validation approach in situations where the reference network is not available. After this evaluation, transcriptomic regulatory relationships between groups can be compared, allowing for the identification of marker arcs. The package also supports the coarse-graining of SCT data to reduce computational costs, probabilistic reasoning, and comprehensive visualizations. A detailed description of all the implementations of *scstruc* is available in the Methods section. We summarize the results from the simulation experiment and from the actual SCT data from mouse and human kidneys to illustrate the typical outputs and insights produced by *scstruc*.

**Figure 1.**
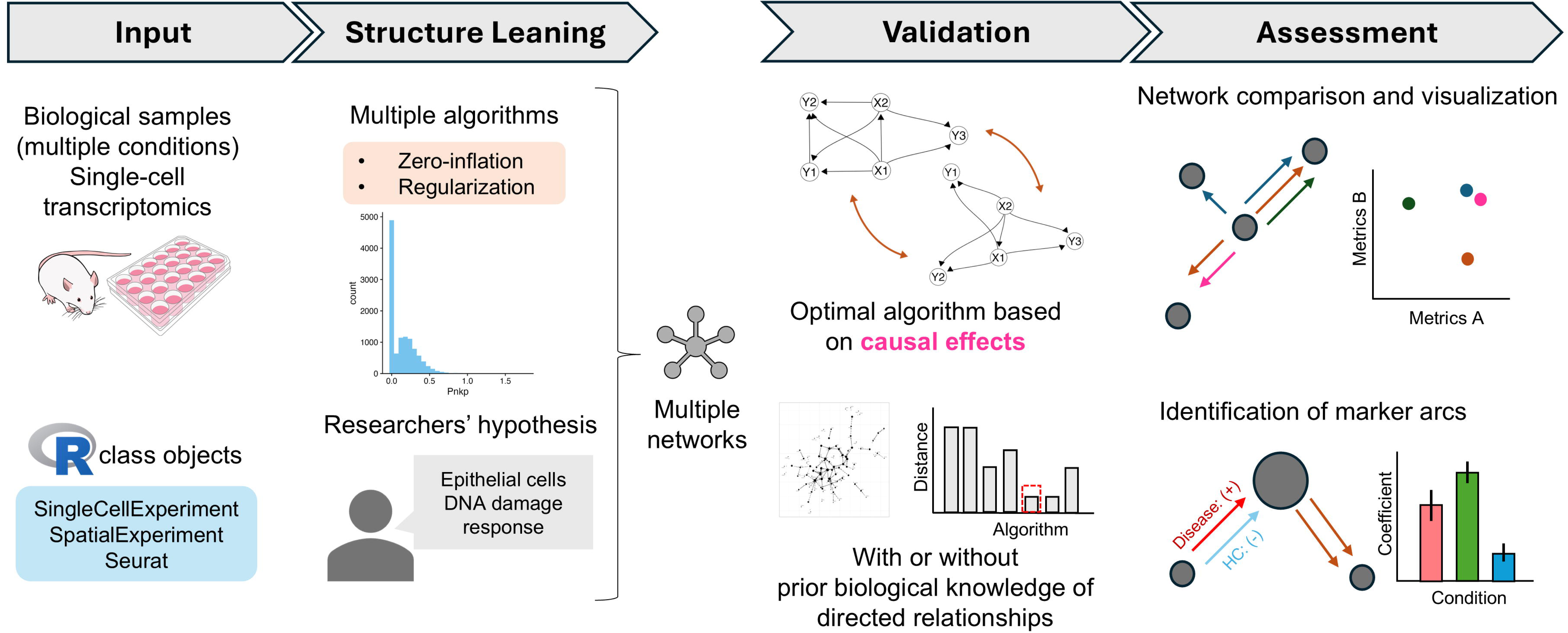
The typical workflow of the package *scstruc.* A typical workflow of the package is shown. Single-cell transcriptomics (SCT) datasets with multiple samples are stored in the R class object and used as input for multiple structure learning algorithms tailored for SCT data. After learning, networks are validated based on metrics including those assessing causal effects between genes regardless of the availability of reference directed networks. Marker arcs differentiating between conditions can be identified based on regulatory relationships in the learned network.

### Algorithm performance in network inference varies by dataset, sample size, and evaluation metric

We first evaluated the performance of implemented structural learning algorithms and commonly used GRN inference software using simulated data sampled from established molecular networks, as well as that simulated by SERGIO (single-cell expression of genes *in silico*). The performance of each GRN inference method was evaluated based on the ground-truth DAGs between genes using implemented metrics, including those assessing causal validity. The results for the 100, 500, and 1000 observations sampled from the two reference molecular networks, consisting of 46 and 107 nodes, and 2700 observations of 100 genes generated by SERGIO, are summarized in Supplementary Table S1.

Specifically, in the ECOLI70 dataset with 100 samples, the hurdle model with the modified Bayesian information criterion (zBIC) achieved the lowest structural hamming distance (SHD) and structural intervention distance (SID). Greedy Equivalence Search (GES) performed best in terms of F1-score. For a sample size of 500, LASSO with cross-validation (CV) outperformed others in SHD, while the hurdle model with zBIC was also strong in SID, and GES had the highest F1-score.

For the larger ARTH150 dataset with a sample size of 100, CCDr achieved the lowest SHD, while the min-max concave penalty (MCP) with CV outperformed in minimizing SID. LASSO with CV showed the highest F1-score in this group. At sample sizes of 1000, the hurdle model with zBIC scoring was the best for SHD, and the Max-Min Hill Climbing (MMHC) algorithm performed well in reducing SID. Smoothly-clipped absolute deviation (SCAD) with CV achieved the highest F1-score.

Generally, the CCDr algorithm was the fastest structure learning algorithm, considering that the algorithm learns the networks for multiple penalty factors. zBIC refers to the scoring function based on the hurdle model, and the details for all the algorithms are provided in the Methods section. The plots for the calculated SHD, SID, and F1-score are shown in Figure 2A. The inferred networks of ECOLI70 and ARTH150, achieving the highest F1-score and the lowest SID, are shown in Figures 2B and 2C, respectively. While both networks aim to reconstruct the same ground-truth system, the results differ substantially in terms of edge selection. The network maximizing F1-score tends to include more edges overall, prioritizing recall, whereas the network minimizing SID results in a sparser structure.

**Figure 2.**
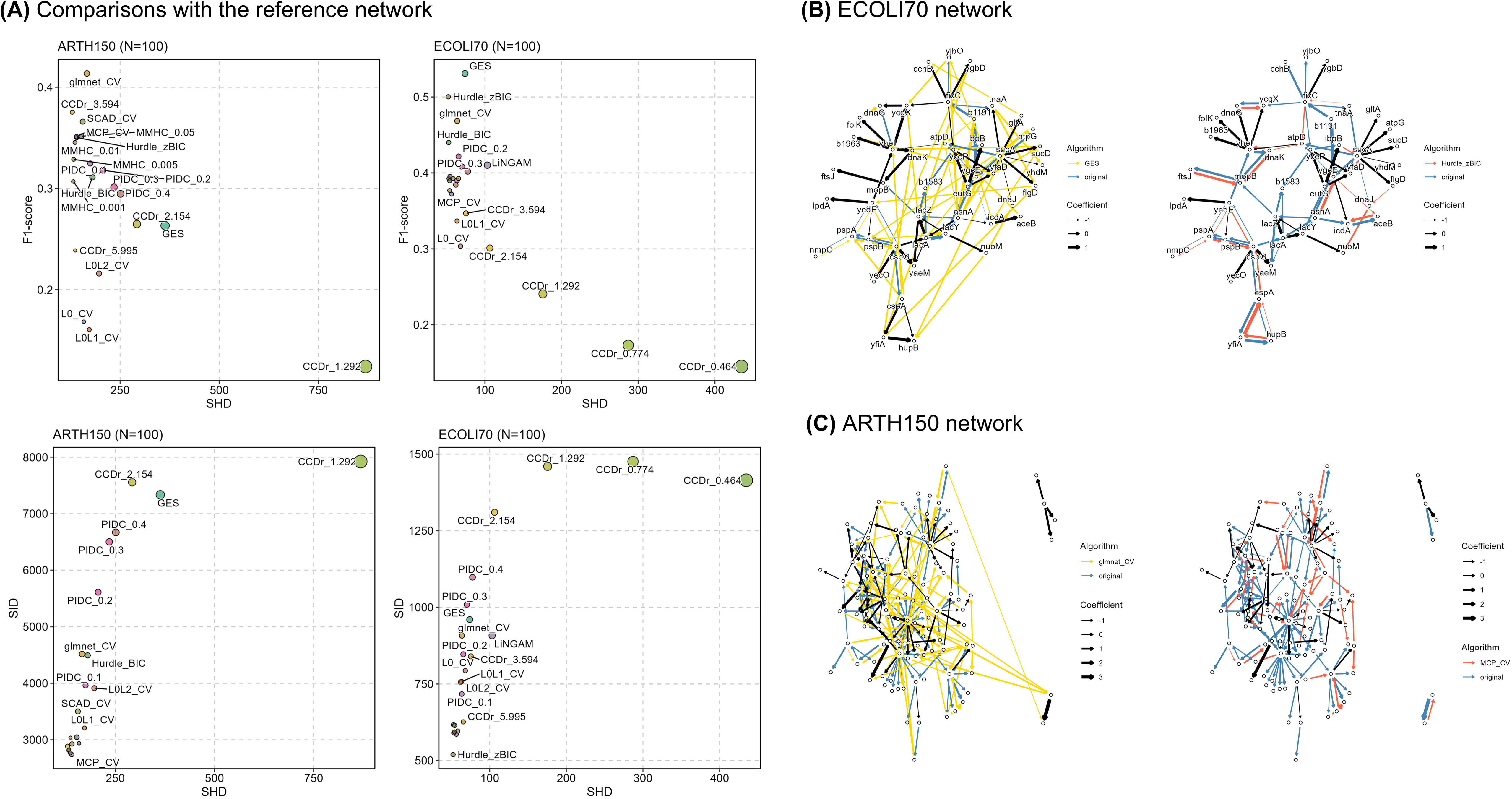
Evaluation of metrics of structure learning algorithms in the simulated dataset. (A) The relationship between structural hamming distance, structural intervention distance, and F1-score in various algorithms. The point size indicates the number of arcs. The CCDr algorithm is labeled with the λ value. MMHC algorithm is labeled with their α values. The results are based on the 100 observations sampled from the reference gene regulatory networks of ECOLI70 and ARTH150 from the *GeneNet* R package. The sampling was performed five times and the average values were plotted. (B) The inferred and original network of ECOLI70 and (C) ARTH150. The yellow arcs are from the network with the highest F1-score, the red arcs with the lowest SID, and the blue are from the original network. True positive arcs are colored black. The arc width corresponds to the fitted parameters. Only the connected nodes are shown here.

For more realistic synthetic data of SCT, network inference from data generated by SERGIO was performed. The results showed lower recovery in all the methods and algorithms compared to the two simulation datasets mentioned above. The higher accuracy was achieved with the clean data than with the data with noise. This result, the introduction of technical noise results in reduced accuracy compared to clean data, is consistent with the findings reported in the original study. Also, the hurdle model performed the best in terms of F1-score in both clean and noisy data. This finding suggests that the hurdle algorithm implemented in *scstruc* effectively handles zero-inflated data. These results are summarized in Supplementary Figure S1.

Overall, the simulation results indicate that the optimal algorithm is dependent on the specific network, the number of genes, and the sample size. These findings highlight the importance of selecting algorithms based on the characteristics of the data and the evaluation metric of interest in each investigation. Importantly, it was also suggested that SID, SHD, and F1-score, which measure causal correctness, structural accuracy, and predictive performance, respectively, have distinct characteristics in simulated data from the reference GRN. Thus, depending on whether we prioritize predictive accuracy or causal interpretability, we may favor very different models.

While *GRNBoost2* and *GENIE3* were included in our comparative analysis, we encountered difficulties in producing DAGs from their outputs. Although both methods are capable of inferring directed relationships between genes and ranking them, they do not explicitly produce DAGs by thresholding by various parameters, which makes direct comparisons with the BN approaches challenging. This limitation highlights the advantage of *scstruc,* which is specifically designed to infer causal relationships by constructing DAGs, thereby providing more interpretable results for causal inference.

Subsequently, we conducted additional experiments validating the Intersection-Validation approach on SHD and SID using the generated data from ECOLI70 and ARTH150 data and presented the results in Supplementary Text S1. The Intersection-Validation approach can rank the algorithms or software based on the specified metrics in situations where the reference network is not available, and the results highlighted that the approach is a useful indicator for SHD and SID as well.

### Biological pathway-specific optimal algorithm selection in the actual SCT dataset

We subsequently inferred and evaluated the network and identified marker arcs using actual SCT datasets obtained from mouse and human kidneys. A mouse dataset includes SCT data from 16 kidney samples, 8 from mice expressing the early diabetic kidney disease (DKD) phenotype, and 8 from controls.^32^ Diabetic nephropathy is a critical comorbidity in diabetic patients affecting various structures in the kidney, especially glomeruli. The samples were processed for the enrichment of glomeruli before sequencing. We identified 24 cell types using Walktrap graph-based clustering.^33^ Structural inference was performed in the mesangial cell type we focused on, as described in the Methods section.

We extracted DAGs from the five biological pathways related to the differentially expressed genes between the conditions and compared the SHD and SID based on the inferred networks. The results of all metrics are presented in Supplementary Table S2. The number of arcs in the extracted reference networks ranges from 21 to 91. The benchmarking results comparing SHD, SID, and F1-score are summarized in Figure 3A. As expected for sparse networks, the SHD decreased with the number of inferred arcs (Figure 3A). The F1-score computed against the validated biological networks was generally low in all the algorithms tested, with the penalty-based regression performing better. Different algorithms demonstrated superior performance between F1-score, SHD, and SID metrics, respectively, suggesting that these measures hold distinct significance in the context of GRNs learned from the actual SCT data, as same as simulation data.

**Figure 3.**
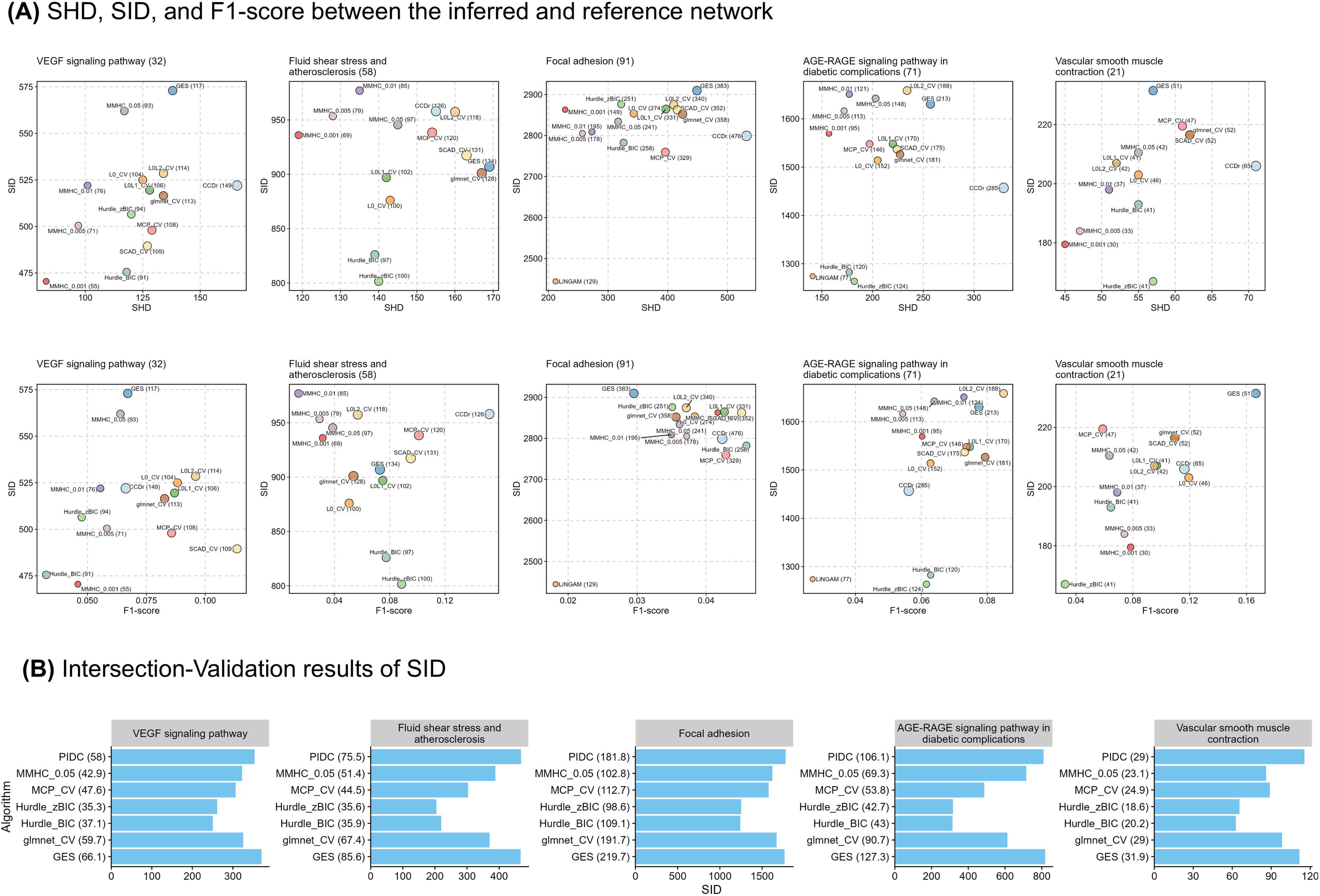
Evaluation of causal relationships in the diabetic kidney disease dataset. (A) The structural hamming distance, F1-score, and structural intervention distance (SID) between the directed acyclic graph (DAG) obtained from KEGG PATHWAY and inferred graphs from each method are summarized. The title indicates the KEGG PATHWAY description with the original node number in the graph. The point indicates the algorithm with the inferred arc numbers. In the Vascular smooth muscle contraction pathway, *PIDC* was omitted from the SID and F1-score plot as the recall and precision were zero. (B) The barplot shows the Intersection-Validation results. The mean SID between inferred networks from subsampled data and the intersection of inferred networks from the original data is shown. The number in the brackets indicates the mean arc number in the network inferred from subsampled data.

In the comparative analysis of algorithm performance based on the SID across different pathways, *PIDC*, the hurdle model with zBIC scoring, and LiNGAM performed best across five pathways. The lower SID indicates a closer alignment with the reference biological networks in terms of causal relationships. The results indicate that selecting the algorithm tailored to that pathway may be more advantageous.

Furthermore, Intersection-Validation results suggest that the hurdle algorithm performs consistently better, particularly in complex pathways such as the AGE-RAGE signaling in diabetic complications and Focal adhesion, achieving the lowest SID values in these cases. These results, summarized in Figure 3B, highlight the effectiveness of the hurdle models, which appear to outperform the alternatives in producing causally plausible biological conclusions.

These findings illustrate how *scstruc* can elucidate pathway-specific behaviour of network inference performance and how important it is when selecting algorithms for directed GRN inference tasks based on multiple metrics.

### Optimal network captures plausible marker arcs

Among the inferred networks, algorithms based on the hurdle model performed better in terms of SID in the AGE-RAGE signaling pathway in diabetic complications. Building on this, we constructed the averaged DAG from 50 bootstrap samples, fitted the per-cell-type parameters of the associated BN, and compared across cases and controls. The reference network and the inferred network in the investigated mesangial cell type are shown in Figures 4A and 4B.

**Figure 4.**
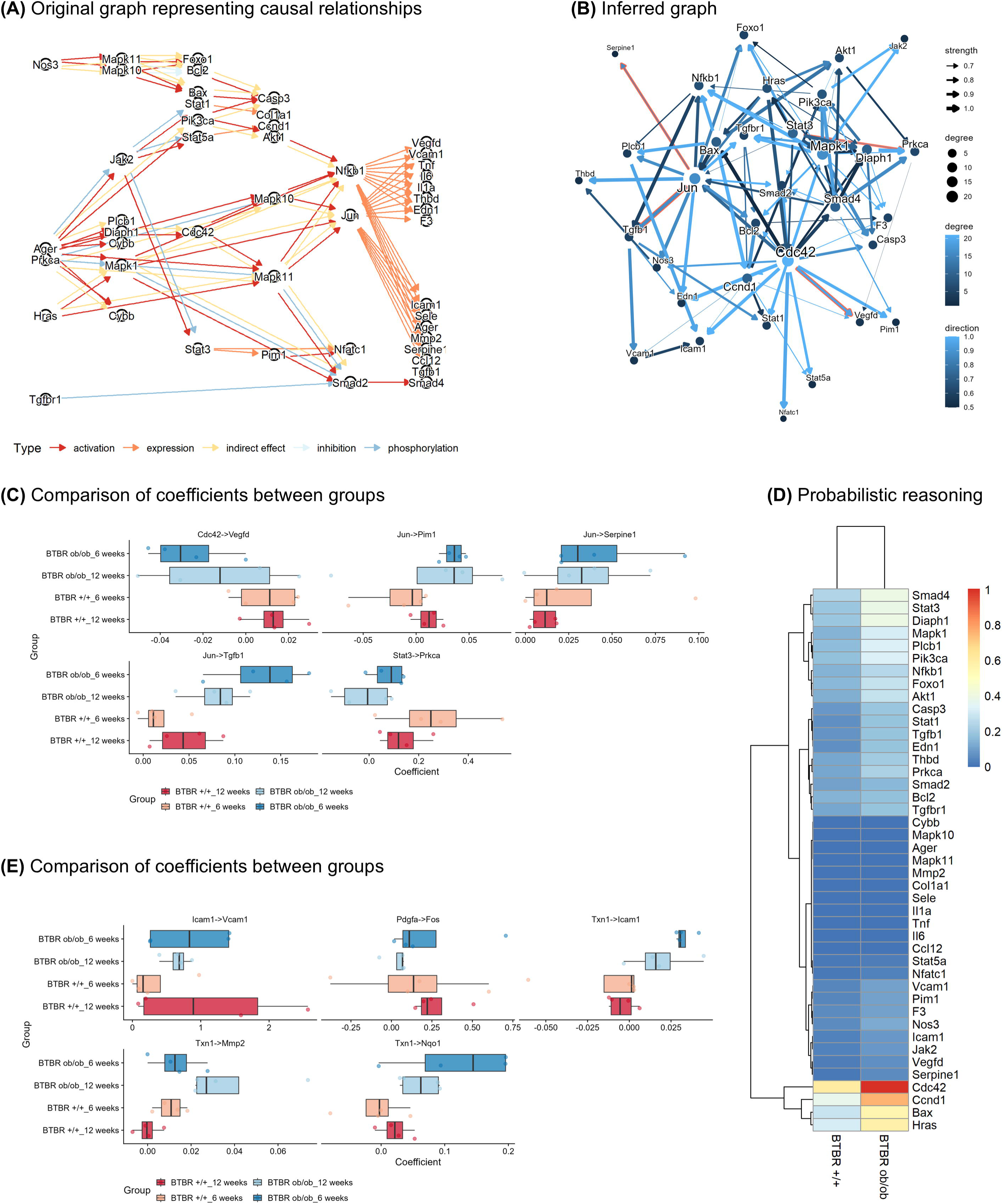
The results of marker arc identification in diabetic kidney dataset. (A) The directed acyclic graph was obtained from the original graph and used to evaluate the causal relationships. (B) The visualization of the inferred graph by the hurdle model based algorithm. The highlighted arcs are those identified as markers distinguishing between diabetic kidney disease and control mice. The nodes are sized based on the degree of the nodes. (C) The boxplot shows the result of the comparison of coefficients in the AGE-RAGE signaling pathway in the diabetic complications pathway. (D) The heatmap shows the results of probabilistic reasoning based on the perturbation of the expression of the gene Jun in the selected network. (E) The result of the comparison of coefficients in Fluid shear stress and atherosclerosis pathway.

Marker arcs mainly originated from the hub genes, such as *Jun* and *Cdc42* in the inferred network. The *Jun* to *Tgfbr1* and *Serpine1* arcs agree with the original KEGG PATHWAY. These relationships were dysregulated in DKD mice (BTBR ob/ob in Figure 4C), which supports the reported pathophysiology in mesangial cells.^34^ We subsequently investigated the inferred networks by probabilistic reasoning. Mean *Jun* expression of each group of mouse strain was used for the evidence and thresholding, and samples above or below the threshold were retained. Mean sampled values across genes are shown in the heatmap (Figure 4D). The differences of *Cdc42 are* more prominent in the mouse diabetic phenotype, suggesting the active regulation of this relationship through *Cdc42* to *Jun* in the mouse with a diabetic phenotype.

We further investigated the fluid shear stress and atherosclerosis pathways using the same method, identifying the marker arcs (Figure 4E). From the results, the coefficients of regulation in the antioxidant genes (*Txn1*, *Nqo1*) were dysregulated. Oxidative stress is reported to play a central role in DKD progression and pathophysiology, and antioxidative stress response has been suggested to occur in mesangial cells.^35^

These dysregulations of regulatory relationships in the mesangial cell types could be linked to pathogenesis in early DKD. The *scstruc* provides all the tools to make these links and makes establishing them straightforward. We additionally examined the SCT dataset of human kidneys with diabetes under the treatment of sodium-glucose transporter 2 (SGLT2) inhibition and confirmed the usefulness of the approach. We presented the results in Supplementary Text S2.

## Limitations of the study

The limitations of *scstruc* were that it uses only quantitative transcript values and does not consider other factors, such as the relationship with the transcription factors. However, inference based on whitelisting these relationships is possible within the function. Also, the evaluation of biological pathways in the paper was not comprehensive but limited to those related to differences in the specific cell types. The analysis of the other cell types across the genes related to the other pathways is needed to verify the approaches further.

## Discussion

We described the R package *scstruc*, which aims to infer directed relationships between genes in SCT data based on experimental hypotheses, modeling and evaluating them as BNs. The package implements inference, validation, and comparison of parameters and network structures between conditions, which we demonstrated on real-world SCT data obtained from mouse and human kidneys.

The package is designed to apply a wide selection of algorithms tailored to account for the unique characteristics of SCT data to infer GRN through BN. It also provides the ability to evaluate the causal validity of these inferred networks both in situations where the reference networks are available or not. Moreover, *scstruc* offers an integrated framework for marker arc detection within these validated networks, significantly enhancing the user’s ability to analyze and interpret SCT data.

Although predictive metrics such as the F1-score, area under receiver operating characteristic curve, and precision recall curve are important and commonly used for evaluating the algorithms for recovering the ground-truth networks, the performance is generally low in SCT datasets.^36–38^ In addition to these metrics, we considered assessing causality critical in the context of GRNs: *scstruc* provides this novel functionality for evaluating causal relationships because they are biologically most relevant for network inference. Also, *scstruc* can assign directions based on scoring algorithms using undirected networks produced by other algorithms, broadening the usage of published GRN inference software.^39^

Metrics assessing directed relationships, one of the characteristics of BN analysis, typically requires access to the reference directed regulatory relationship. However, the number of databases describing the regulatory relationships of whole genes as directed or DAG is limited,^40^ and no established data or database currently exists. Therefore, *scstruc* can use either the Intersection-Validation approach or DAGs obtained from the KEGG PATHWAY database, which contains manually curated regulatory relationships between genes for a variety of functional categories, as an alternative in this type of evaluation.

BN structure learning has been applied to GRN inference in past literature. In this study, we evaluated the application of various algorithms such as regularization-based regression in BN structure learning from SCT data. While regularization-based learning showed usefulness in recovering true positive arcs compared to the other methods in simulation experiments, the evaluation results based on real-world SCT data indicated that the recovery of arcs aligned with the network obtained from biological databases was low depending on the subset genes and cells. Importantly, the structure learning based on hurdle models or *PIDC*, designed for the SCT data, was valuable compared to the alternatives when learning from simulated or actual SCT data. While it was anticipated that inference methods tailored to SCT data would be advantageous, it has been demonstrated that such methods indeed recover the biologically meaningful arcs with regard to causal relationships. When performing probabilistic reasoning or similar analyses based on networks derived from SCT data, it is desirable for the network to be as close as possible to a biologically validated network in terms of causality. This package greatly simplifies the evaluation for such purposes.

In summary, our *scstruc* package brings BN structure learning and evaluation of causal validity to SCT data, allowing for easy and efficient hypothesis generation. The integrated approach of including network inference, benchmarking, and comparison in a single package makes modeling and interpreting SCT data based on the user’s experimental hypotheses straightforward, providing a coherent workflow from structure learning to biological interpretation. Furthermore, *scstruc* goes beyond probabilistic modeling to provide many of the tools needed for causal discovery and inferences to further advance the user’s ability to draw clinically or biologically relevant insights.

## Resource availability

### Lead contact

- Requests for further information and resources should be directed to and will be fulfilled by the lead contact, Seiya Imoto.

### Materials availability

- This study did not generate new unique reagents.

### Data and Code availability

- This paper analyzes existing, publicly available data, accessible at Gene Expression Omnibus under the accession identifiers GSE218563 and GSE220939.
- All original code has been deposited at https://github.com/noriakis/scstruc.
- Any additional information required to reanalyze the data reported in this paper is available from the lead contact upon request.

## Acknowledgements

This study is partially supported by Takeda Science Foundation, JSPS KAKENHI 25K21331; and grant K25-2170 from the International Joint Usage/Research Center, the Institute of Medical Science, the University of Tokyo.

## Author contributions

Noriaki Sato: Conceptualization, Data curation, Formal analysis, Funding acquisition, Investigation, Methodology, Project administration, Software, Resources, Visualization, Writing – original draft, Writing – review & editing. Marco Scutari: Conceptualization, Investigation, Methodology, Software, Writing – review & editing. Seiya Imoto: Investigation, Methodology, Project administration, Writing – review & editing.

## Supplementary Information

SupplementaryFigureAndText.docx.

Supplementary Figure S1. The performance metrics of SERGIO generated data.

Supplementary Text S1. Intersection-Validation approach on the SHD and SID.

Supplementary Text S2. The analysis of the kidney human biopsy dataset.

SupplementaryTables.xlsx.

Supplementary Table S1. The comparison between the reference and inferred network for simulated data.

Supplementary Table S2. The comparison between the reference and inferred network for actual data.

## Methods

### Software architecture and Implementation

The package code is available at https://github.com/noriakis/scstruc. The package is structured into modules which provide all the functionality required for learning GRNs based on and evaluating BNs from SCT data. Users can directly pass *Seurat*, *SingleCellExperiment*, or *SpatialExperiment* objects, the standard class objects to encode SCT data in R, to the functions in *scstruc*. The package modules can then perform structure learning, evaluate and validate the inferred networks, and compare the directed relationship between groups of interest by leveraging *bnlearn* and other R packages.^41^

BNs are a class of graphical models defined over a set of random variables *X* = {*X*_1_, …, *X_N_*}, which are associated with the nodes in a DAG *G*. The graphical separation between nodes in *G* implies the conditional independence of the corresponding variables and leads to factorization:

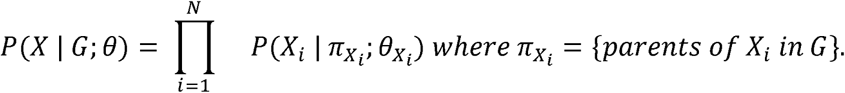

In GRNs, each node corresponds to a gene, and we try to learn the graphical structure of the network (that is, the arcs that appear in it) using the per-cell gene expression matrix.

### Learning the structure of GRNs

The *scstruc* package allows for learning BNs from SCT data with a broad selection of algorithms, including both two-stage (or hybrid) and one-stage approaches. The former restricts each node’s candidate parents and children and then learns the structure that maximizes the score under the resulting constraints.^42^ The latter learns the parents of the nodes directly from the data. For instance, the CCDr algorithm combines a MCP and block coordinate descent to learn the BN.^43^

The *scstruc* package implements several two-stage approaches: the L1-regularized Markov Blanket,^44^ L0-regularized regression, MCP, and the SCAD penalty. The penalty factor (usually denoted with λ) can be chosen based on the CV or the lowest BIC. In addition, it provides wrappers around the structure learning algorithms available from *bnlearn* and the CCDr algorithm implemented in the *sparsebn* package.^45^

To handle possible zero inflation, *scstruc* uses the multivariate hurdle model, a hierarchical model that handles the excess zeroes separately from other observations. It first learns the edges in this model using *HurdleNormal* and grouped lasso,^23^ which uniquely implements a multivariate hurdle model with a multivariate normal distribution. As the edges are undirected, *scstruc* learns the structure of a BN by blacklisting all other possible edges in the HC or GES algorithm based on score maximization. The scoring function in HC can be chosen from the default score implemented in *bnlearn* or those from the hurdle model. This corresponds to the sum of the BIC values of the logistic regression model indicating whether the gene expression is zero and of the Gaussian regression model for continuous values (denoted as zBIC). Let *Y* = [*y_ij_*] denote the log-normalized expression value of the gene *i* in subset cell *Z* and [*z_ij_*] a 0-1 indicator value of whether the gene expression is zero. *scstruc* fits the model using the identified neighbour gene set *G* and calculates the score as follows:

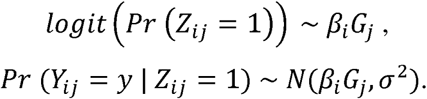

The sign of the BIC score is reversed in keeping with the convention that the optimal structure maximizes the scoring function. We chose the *MAST* package for scoring the hurdle model because it uses *bayesglm* in *arm* internally in hurdle modeling,^46^ and therefore has robust performance in differential expression analysis. Finally, *scstruc* improves the BN’s robustness by averaging the network structures learned from bootstrap samples and evaluating the strength and direction confidence of the inferred edges.^47^

Since learning network structures from all the genes on a standard SCT platform (typically more than 10,000) is not feasible in many computational environments, *scstruc* fetches data from biological pathway databases such as the Gene Ontology and Kyoto Encyclopedia of Genes and Genomes (KEGG) and subsets the genes in the pathway for the inference.^48,49^ Thus, the user can choose custom sets of input genes to match their research purposes or hypotheses.

### Preprocessing the SCT data

SCT data are initially cleaned using standard quality control steps: filtering low-quality genes and cells and transforming counts to the logarithmic scale. The resulting class objects storing the pre-processed expression matrix, such as *SingleCellExperiment* or *SpatialExperiment,* can then be used safely in *scstruc*. *Seurat* class objects can be converted beforehand using the appropriate method from that package.

Processing multi-sample SCT data sets poses a substantial computational burden because they typically contain millions of cells. The *scstruc* package solves this issue by merging cells with highly similar transcriptomic profiles into metacells using the *SuperCell* R package before structure learning.^50^

### Fitting the parameters, identifying the markers, and visualizing results

After learning the structure of the network, *scstruc* can learn its parameters, the regression coefficients associated with the parents of each gene, from the gene expression matrix of each cell type per sample. The package identifies the important regulatory relationships that differentiate two conditions using a feature selection algorithm. The default choice is *Boruta* from the CRAN package of the same name, which performs feature selection using random forests with shadow variables.^51^ Other methods for feature ranking, such as importance estimates from *xgboost,* are also supported.^52^

An additional key feature of *scstruc* is its visualization options, which allow users to interpret the network structure intuitively. The package supports displaying arc strengths, highlighting marker arcs, plotting subnetworks, and facilitating comparisons with reference networks, similar to those that can be achieved by Cytoscape.^53^ *ggraph* was used internally for network plotting.

### Evaluating the causal relationships in the learned networks

We conducted a simulation experiment to showcase how to use *scstruc* to compare various metrics and algorithms. We chose the Gaussian BNS, ECOLI70 and ARTH150, from the *GeneNet* package as reference networks and sampled data sets with 100, 500, and 1000 observations from them.^54^ This reflects the typical metacell numbers in the cell cluster in analyses in the literature. Also, SERGIO is employed to simulate single-cell gene expression data with technical noise.^38^ We used the DS1 dataset consisting of 100 genes and 2700 cells, generated from the network with 258 edges. The clean expression matrix as well as that with the addition of outlier genes, library size effects, and dropouts (percentile parameter of 65), were used as input.

> We considered the following algorithms for comparison.

● The L0, L0L1, and L0L2 regularization from the *L0Learn* package, with the default algorithm of coordinate descent.^55,56^ L0L1 and L0L2 combine the L0 norm with L1 and L2 norms.
● The LASSO from the *glmnet* package.^57^
● MCP and SCAD from the *ncvreg* package.^58^
● The HC and MMHC algorithms from the *bnlearn* package. MMHC used significance thresholds 0.001, 0.005, 0.01, and 0.05.
● The CCDr algorithm with the default settings (λ=20, α=100, and γ=2) to match MCP.
● LiNGAM and GES from the *pcalg* package.^59^
● The hurdle model.

We compute several metrics from the learned networks to assess the accuracy of the structure learning algorithms. *scstruc* implements arc number; true positive, false negative, and false positive arcs; true positive rate (TPR), precision and recall; F1-score and SHD; BIC and the Kullback-Leibler (KL) divergence from the reference network.

Molecular networks are often used for hypothesis generation through interventions, so metrics describing how accurately they represent causal pathways are very relevant. For this reason, we also considered the SID.^29^ SID compares two DAGs based on their causal inference statements and measures the number of false intervention distributions. We used the symmetrized version of SID implemented in *bnlearn* or the CRAN package *SID*.

Most metrics above evaluate the outputs of structure learning algorithms against the true network for the data, which is often not readily available when we work with experimental SCT data. To simulate this scenario, we extract the directed relationships between the genes of interest from the molecular pathway database, KEGG, and compare the inferred networks with the resulting DAG. We construct a DAG from the set of directed relationships by first identifying them in the KEGG PATHWAY based on gene-to-gene interactions, then isolating the largest components and removing the feedback arc set in the component. In this way, we extract a DAG from each pathway, and we can use inference on its genes to evaluate the learned networks. A feedback arc set is a subset of edges in the graph whose removal breaks all cycles in the graph.

Finally, we also evaluated Intersection-Validation to select the algorithms where the biological reference network is not available.^60^ This approach considers the intersection of networks learned from multiple algorithms. It calculates metrics comparing the intersection network and the networks learned from subsamples of the original data by each algorithm. The original paper considered the partial hamming distance as the main metric, which proved to be a good substitute for the comparison with the reference network. We ranked the algorithms using SID, a sample size of 250, and 10 replicates in the following.

### Comparison with other GRN inference software and algorithms

In addition to various learning algorithms, we compared the directed graphs produced by other software. *GRNBoost2* and *GENIE3* provide directed relationships between genes based on per-gene regressions and rank them.^26,61^ We included the inferred networks thresholded by various parameters only if the network is directed and acyclic. The *PIDC* algorithm uses a partial information decomposition to output an undirected network, and we determined the edge orientation with HC and BIC.^39^ The GES can be additionally used.

### Data set description

We also showcase using *scstruc* with real-world mouse data on early diabetic kidney disease (DKD). This data set includes SCT data from 16 kidney samples, 8 from mice expressing the diabetic phenotype, and 8 from controls.^32^ After quality control, we converted the log-transformed count matrix into a representative metacell count matrix and applied batch correction using the *fastMNN* function in *batchelor*.^62^ We identified cell types using the *EasyCellType* package to perform a gene set enrichment analysis on the marker gene scores identified by the uncorrected expression matrix.^63^

The uncorrected log-count matrix of the metacells per cell type was used for the input of the analysis, as a gene-based comparison could be biased by batch correction. We learned the network structures from them with the algorithms in *scstruc* and compared them as described above. We then fitted the parameters of the network with the best performance using the uncorrected log-count matrix.

Subsequently, we identified the regulatory relationships differing between cases and controls for the annotated cell types. We confirmed the findings in Liu *et al*.: several pathways were dysregulated in the glomerular cells of the early DKD mice. We investigated five pathways in KEGG PATHWAY for inference in the cell cluster identified as mesangial cells, including vascular smooth muscular cell contraction, AGE-RAGE signaling pathway in diabetic complications, VEGF signaling pathway, Fluid shear stress and atherosclerosis, and Focal adhesion.

We also analyzed the SCT dataset from a human kidney biopsy specimen under SGLT2 inhibition consisting of 22 samples using the same approach.^2^

### Statistical analysis

Throughout the manuscript, single-cell transcriptomic data were processed using the *scran*, *scater*, *scuttle* R packages, and the *tidyomics* ecosystem based on the *SingleCellExperiment* class object.^64,65^ The graph manipulation and analysis were conducted using *igraph* and *tidygraph*.^66^ The R version 4.4.1 was used.

