## Supplementary Figure and Text for "Causal assessment of gene regulatory network in single-cell transcriptomics data based on Bayesian networks"

**Supplementary Figure S1. The summary of metrics in the experiment using SERGIO**

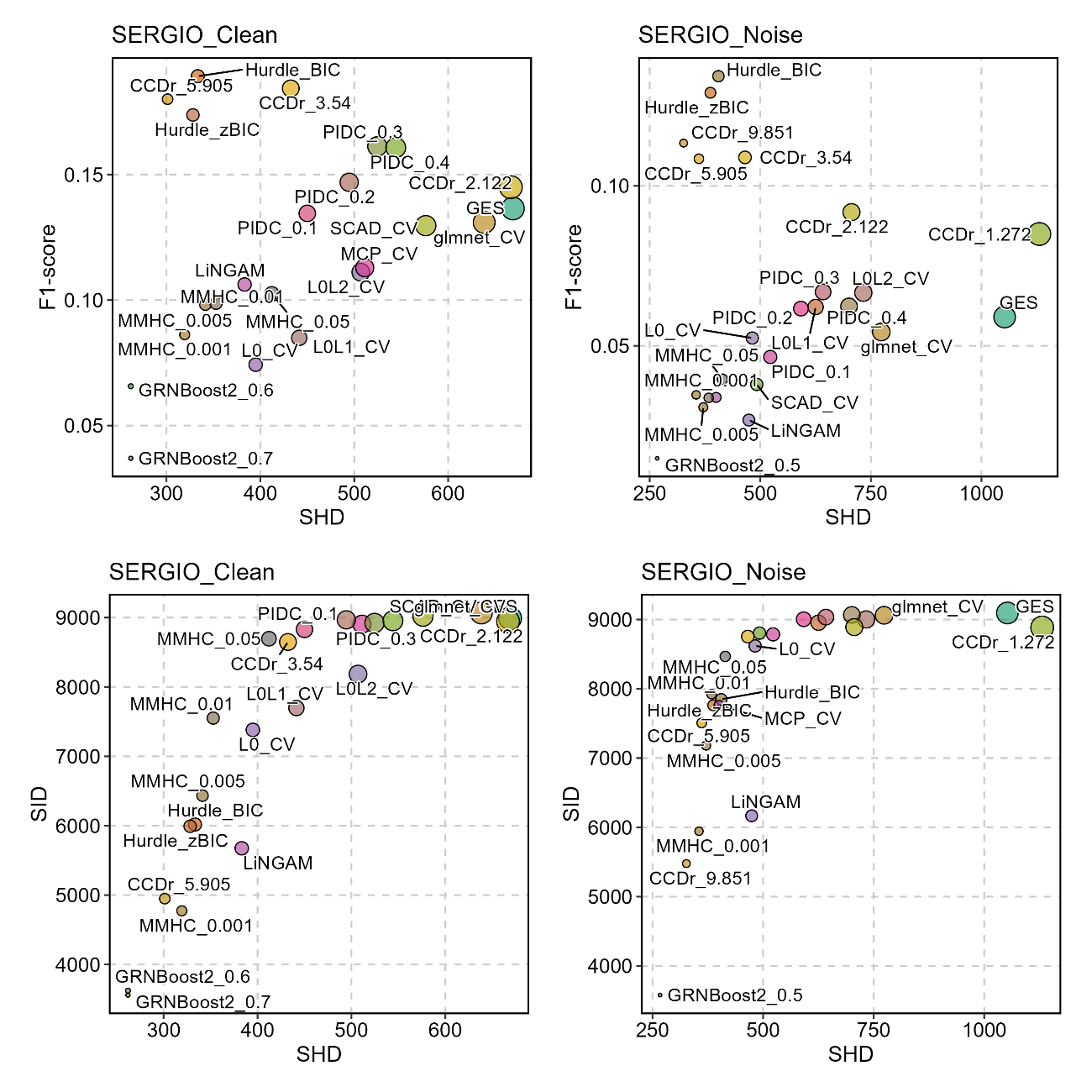

The relationships between SHD, SID, and F1-score are shown. The point color indicates algorithms, and the point size indicates the edge number. The average values obtained from five simulation results are plotted. SERGIO_Clean indicates the use of simulated clean expression data, and SERGIO_Noise indicates the use of simulated count data with the outlier genes, library size, and dropouts. The algorithms without true positive arcs are omitted from the plots.

**Supplementary Text S1. Intersection-Validation approach on the SHD and SID**

We tested Intersection-Validation on structural hamming distance (SHD) and structural intervention distance (SID) using the data generated from ECOLI70 and ARTH150 data.^1^ The symmetric version of SID was used as the metrics. The intersection point was set to 1600, and Hill-Climbing algorithm with the score AIC, BIC, BGE, MMHC, GS, H2PC, and TABU algorithms were used as the candidate algorithms. The intersection of networks inferred from these algorithms was defined as *A0*. Subsampling was performed ten times for sample sizes 50, 100, 200, 300, 500, and 1000 from the original generated data. Mean SHD and SID values between *A0* and inferred network per subsample were obtained. The Pearson’s correlation value between the SHD and SID of the original network and inferred network, and those obtained from subsampling were calculated. The results were summarized in the plot. The edge number of the *A0* was 25 for ECOLI70 and 43 for ARTH150. The results indicate that for both SHD and SID, the correlation of the metrics was high in the larger sample numbers, with better results obtained in SHD.

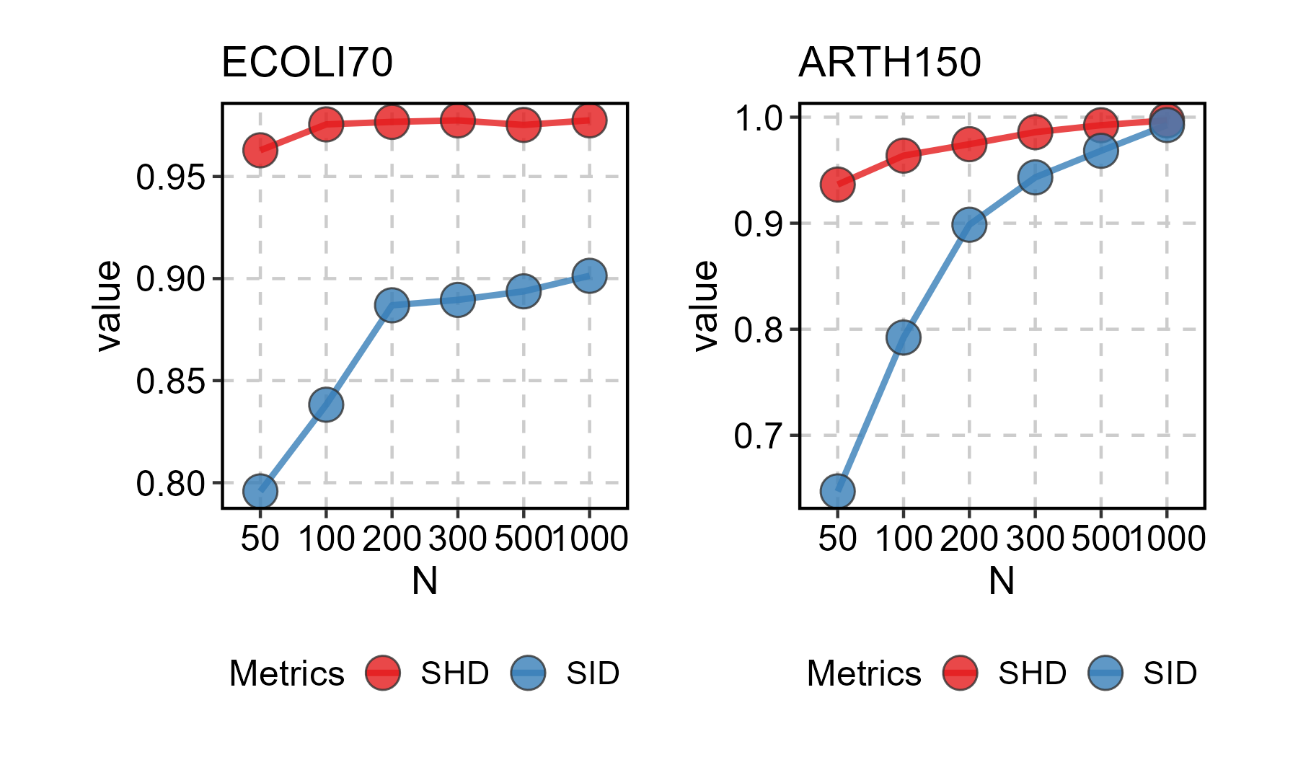

**Supplementary Text S2. The analysis of the kidney human biopsy dataset**

We additionally analyzed the dataset investigating transcriptomic changes in human kidneys under sodium-glucose co-transport 2 (SGLT2) inhibition using the single-cell transcriptomic data (accession ID: GSE220939).^2^ The dataset consists of 22 samples of kidney biopsy specimens with or without diabetes under treatment of SGLT2 inhibition or not. The same analysis method as the mouse kidney dataset was applied. The cell type was annotated based on

As the original manuscript focused on the transcriptomic changes of the mTORC1-signaling pathway, we inferred the network of genes involved in the corresponding pathway (hsa04150). We selected proximal tubular cells for the investigation as the original study reported the alterations of transcripts. We inferred the network using multiple algorithms and found the best network in terms of minimizing the structural intervention distance between the directed acyclic graph originating from the original pathway.

The results indicated that the hurdle model with BIC scoring achieved the best performance. We identified the arcs distinguishing between samples under SGLT2 inhibition and not based on the feature importance from *xgboost* function, and we plot the corresponding inferred network and identified marker arcs below.

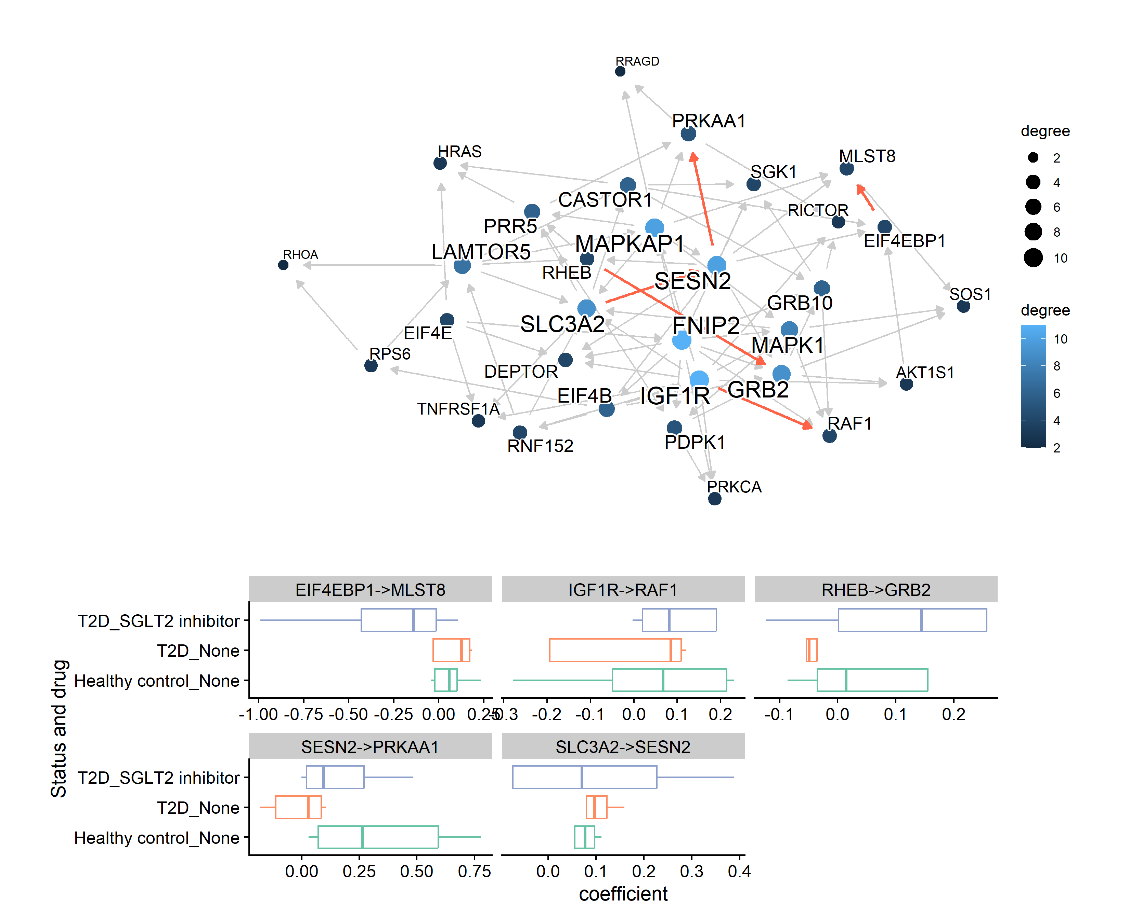
The results suggest that SGLT2 inhibitor treatment normalizes changes caused by type 2 diabetes (T2D) toward a healthy control state. This finding supports the original study's conclusion that SGLT2 inhibitors normalize transcriptomic changes induced by T2D and further demonstrates that the regulatory relationships within the network undergo similar changes.

By utilizing this package, not only can individual transcriptomic changes be analyzed, but also the regulatory relationships within an estimated network that more closely resembles a biologically validated network. This approach enables a deeper understanding of SCT data.
